# Extraction of saccades from eye movements triggered by reflex blinks

**DOI:** 10.1101/132605

**Authors:** Uday K. Jagadisan, Neeraj J. Gandhi

## Abstract

The trigeminal blink reflex can be evoked by delivering an air puff to the eye. If timed appropriately, e.g., during motor preparation, the small, loopy blink-related eye movement (BREM) associated with eyelid closure disinhibits the saccadic system and reduces the reaction time of planned eye movements. The BREM and intended eye movement overlap temporally, thus a mathematical formulation is required to objectively extract saccade features – onset time and velocity profile – from the combined movement. While it has been assumed that the interactions are nonlinear, we show that blink-triggered movements can be modeled as a linear combination of a typical BREM and a normal saccade, crucially, with an imposed delay between the two components. Saccades reconstructed with this approach are largely similar to control movements in their temporal and spatial profiles. Furthermore, activity profiles of saccade-related bursts in superior colliculus neurons for the recovered saccades closely match those for normal saccades. Thus, blink perturbations, if properly accounted for, offer a non-invasive tool to probe the behavioral and neural signatures of sensory-to-motor transformations.

**New and noteworthy:** The trigeminal blink reflex is a brief noninvasive perturbation that disinhibits the saccadic system and provides a behavioral readout of the latent motor preparation process. The saccade, however, is combined with a loopy blink related eye movement. Here, we provide a mathematical formulation to extract the saccade from the combined movement. Thus, blink perturbations, when properly accounted for, offer a non-invasive tool to probe the behavioral and neural signatures of sensory-to-motor transformations.

## Introduction

As a general rule, an experimental result has a deeper impact when its correlational outcome is fortified by causal evidence. In systems neuroscience, this latter objective is typically addressed by observing how a normal circuit or system responds to perturbations. Apart from supplementing correlational studies, perturbations can also impose new regimes on a system and reveal novel properties that are hidden from observation under normal circumstances. Standard forms of perturbations in nonhuman primate research are electrical microstimulation or chemical injection to excite or suppress activity in a region of neural tissue. While informative, their major disadvantage (at the moment) is the inability to record neural activity at the same site of experimental manipulation. Furthermore, microstimulation induces extraneous activity that could confound interpretation, and neurons take a relatively long time to recover from chemical inactivation which limits their reversibility and application in randomized paradigms. A newer perturbation tool, optogenetics, circumvents some of these limitations, but it has yet to deliver efficient outcomes in higher order mammals.

Our vast knowledge of the neural basis of motor control, sensorimotor integration and cognition has emerged from experimental manipulations applied to the saccadic system. For example, saccades interrupted by microstimulation of the brainstem omnipause neurons (OPNs) have been crucial in demonstrating an internal feedback control mechanism that preserves accuracy (Keller et al. 1996), and bias in target selection after inactivation of the superior colliculus (SC) has highlighted its role in cognitive processing, not just motor control (Gandhi and Katnani 2011; Krauzlis et al. 2013). We have been using an additional, under-appreciated perturbation tool – the trigeminal blink reflex – to obtain behavioral and neural signatures of the (normally hidden) process underlying movement preparation. For example, appropriately timed reflex blinks reduce saccade reaction times significantly (Gandhi and Bonadonna 2005), an observation that we have used to study the link between target selection and movement preparation in a visual search task (Katnani and Gandhi 2013). In addition to providing temporal specificity (like microstimulation) and being non-invasive, it can be combined with measurement of neural activity from desired neural structures. Indeed, it has been used to uncover latent sensorimotor processes (Jagadisan and Gandhi 2016) and assess algorithms for decoding population activity in the SC (Goossens and Van Opstal 2000b).

Blinks remove inhibition on the saccadic system by shutting down the OPNs – these neurons are tonically active when the eyes are stable and turn off during eye movements, thus gating them (Cohen and Henn 1972; Keller 1974; Luschei and Fuchs 1972). The OPNs turn off during blinks (Schultz et al. 2010) due to the fact that the blink induces a slow, loopy, blink-related eye movement (BREM) (Rottach et al. 1998), and we believe the OPN inhibition allows a saccade to follow and overlap with the BREM (Gandhi and Bonadonna 2005). We refer to such movements as “blink-triggered” saccades. Figure 1a shows examples of BREMs evoked during fixation of a central target. Note that the excursion of the eye is significantly large (˜5-10 degrees), although this can vary depending on the subject or initial eye position (Rottach et al. 1998). We therefore expect reduced latency saccades triggered by reflex blinks to be contaminated by the presence of the BREM. Figure 1b shows examples of normal saccades and blink-triggered movements – note that the spatial and temporal profiles of the two sets of movements are different. The initial phase of blink-triggered movements typically overlaps with the initial phase of BREMs (see overlaid purple traces in Figure 1b).

**Figure 1.**
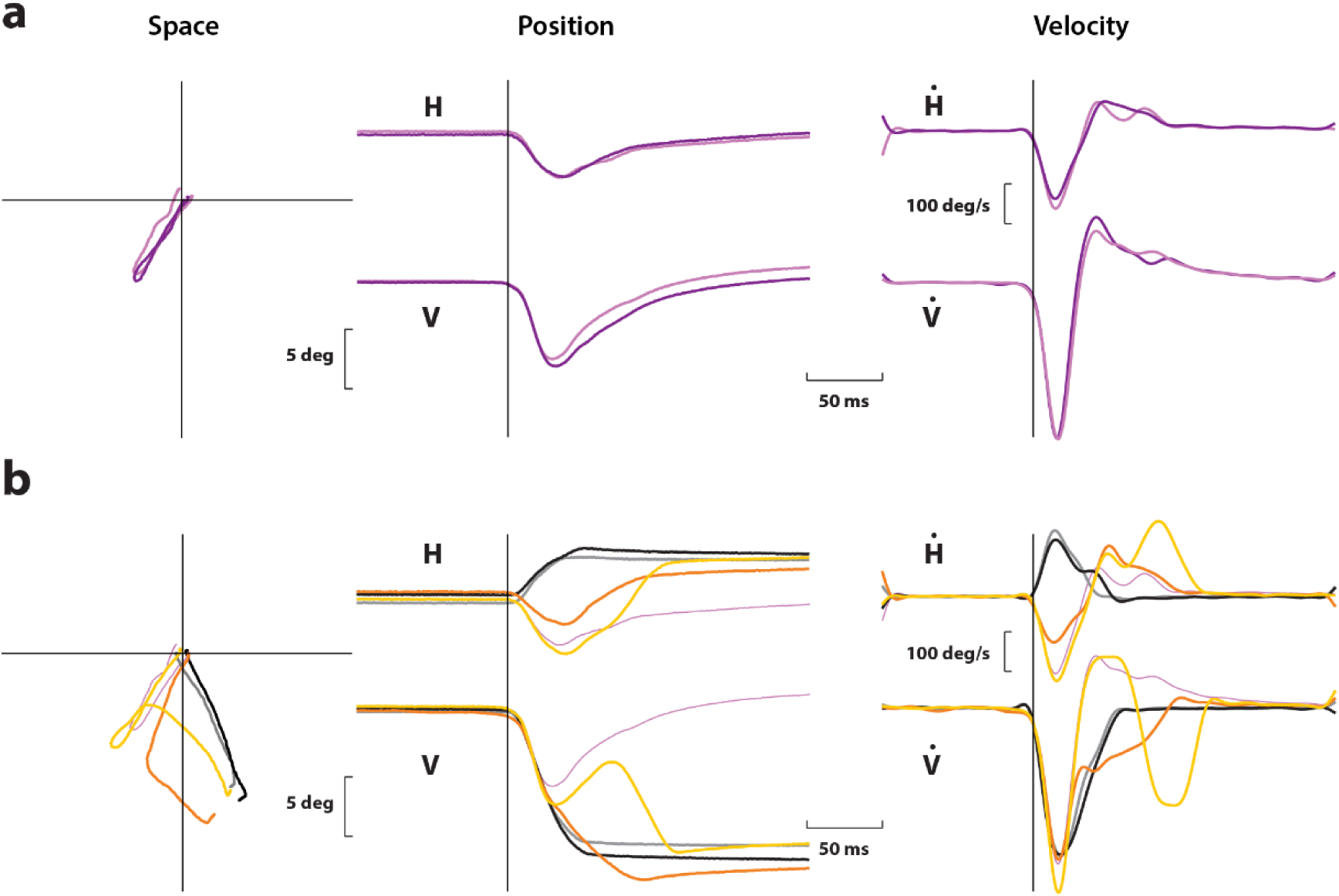
**a**. Example spatial (left), position (middle), and velocity (right) profiles of two BREMs evoked during fixation. The two rows in the middle and right columns show horizontal and vertical signals, respectively. **b**. Examples of two normal saccades (gray and black) and two blink-triggered gaze shifts (orange and yellow) made to the same target. The layout is the same as in **a**. Note the heterogeneity in the profiles of blink-triggered movements. One of the BREMs (thin purple traces) from **a** is overlaid for comparison. Note the similarity between the initial phases of blink-triggered movements and the BREM.

In order to use blinks to study temporal features of movement preparation, it is important to know when exactly during the course of the blink-triggered movement the actual saccade is initiated. A previous study has shown that when a reflex blink is evoked just around saccade onset, the resultant eye movement is slower and does not follow typical saccade kinematics. Such “blink-perturbed” saccades cannot be modeled as a linear combination of the BREM and a control saccade directed to the same stimulus (Goossens and Van Opstal 2000a). Here, we show that blink-triggered movements can in fact be modeled as a linear combination of a typical BREM and a typical saccade, crucially, with an imposed delay between the two components. Saccades reconstructed from the blink-triggered movement using this approach were largely similar to normal saccades in their spatial and temporal dynamics. We further show that saccade-related bursts in SC for the recovered saccade closely resemble those for normal saccades. We compare our results against the previously used velocity threshold-based criterion (Gandhi and Bonadonna 2005; Goossens and Van Opstal 2000a). Our results therefore support the notion that blink perturbations, particularly those that occur well before saccade onset, if properly accounted for, offer a non-invasive tool to probe the behavioral and neural signatures of sensory-to-motor transformations.

## Materials and Methods

### General and surgical procedures

All experimental and surgical procedures were approved by the Institutional Animal Care and Use Committee at the University of Pittsburgh and were in compliance with the US Public Health Service policy on the humane care and use of laboratory animals. We used two adult rhesus monkeys (*Macaca mulatta*, 1 male and 1 female, ages 8 and 10, respectively) for our experiments. Under isoflurane anesthesia, a craniotomy that allowed access to the SC was performed and a recording chamber was secured to the skull over the craniotomy. In addition, a post for head restraint and a scleral search coil to track gaze shifts were implanted. Post-recovery, the animal was trained to perform standard eye movement tasks for a liquid reward.

### Visual stimuli and behavior

Visual stimuli were displayed by back-projection onto a hemispherical dome. Stimuli were white squares on a dark grey background, 4x4 pixels in size and subtended approximately 0.5° of visual angle. Eye position was recorded using the scleral search coil technique, sampled at 1 kHz. Stimulus presentation and the animal’s behavior were under real-time control with a LabVIEW-based controller interface (Bryant and Gandhi 2005). After initial training and acclimatization, the monkeys were trained to perform a delayed saccade task. The subject was required to initiate the trial by acquiring fixation on a central fixation target. Next, a target appeared in the periphery but the fixation point remained illuminated for a variable 500-1200 ms, and the animal was required to delay saccade onset until the fixation point was extinguished (GO cue). Control trials in which fixation was broken before peripheral target onset were removed from further analyses. The animals performed the task correctly on >95% of the trials.

### Induction of reflex blinks

On a small percentage of trials (˜15-20%), we delivered an air puff to the animal’s eye to invoke the trigeminal blink reflex. Compressed air was fed through a pressure valve and air flow was monitored with a flow meter. To record blinks, we taped a small Teflon-coated stainless steel coil (similar to the ones used for eye tracking, but smaller in coil diameter) to the top of the eyelid. The air pressure was titrated during each session to evoke a single blink. Trials in which the animal blinked excessively or did not blink were aborted and/or excluded from further analyses. To obtain blink-triggered movements to the peripheral target, we sought to evoke blinks 100-250 ms after the GO cue, during the early phase of the typical saccade reaction time. In our experimental setup, blink onset occurs approximately 150 ms after the air puff reaches the eye. Thus, air puffs were administered 50 ms before to 100 ms after the GO cue. Trials in which a saccade did not accompany such blinks (i.e., where the gaze did not end up at the target) were removed from further analysis. To obtain BREMs without an accompanying saccade, air puff delivery was timed to evoke blinks during fixation of the central target, 400-100 ms before the onset of the peripheral target. The window constraints for gaze were relaxed for a period of 200-500 ms following delivery of the air puff to ensure that the excursion of the BREM did not lead to an aborted trial.

### Electrophysiology

During each recording session, a tungsten microelectrode was lowered into the SC chamber using a hydraulic microdrive. Neural activity was amplified and band-pass filtered between 200 Hz and 5 kHz and fed to a digital oscilloscope for visualization and spike discrimination. A window discriminator was used to threshold and trigger spikes online, and the corresponding spike times were recorded. The location of the electrode in the SC was confirmed by the presence of visual and movement-related activity as well as the ability to evoke fixed vector saccadic eye movements at low stimulation currents (20-40 μA, 400 Hz, 100 ms). Before beginning data collection for a given neuron, its response field was roughly estimated. During data collection, the saccade target was placed either in the neuron’s response field or at the diametrically opposite location (reflected across both axes) in a randomly interleaved manner. Additional target locations were included in some occasions, particularly when neural activity was not recorded.

### Data analysis and pre-processing

Data were analyzed using a combination of in-house software and Matlab. Eye position signals were smoothed with a phase-neutral filter and differentiated to obtain velocity traces. Normal saccades, BREMs, and blink-triggered eye movements were detected using standard onset and offset velocity criteria (50 deg/s and 30 deg/s, respectively). Onsets and offsets were detected separately for horizontal and vertical components of the movements and the minimum (maximum) of the two values was taken to be the actual onset (offset).

Raw spike density waveforms were computed for each neuron and each trial by convolving the spike trains with a Gaussian kernel (width = 4 ms). For a given neuron and target location, spike densities were averaged across trials after aligning to saccade onset. We also normalized the trial-averaged spike density of each neuron to enable meaningful averaging across the population. The activity of each neuron was normalized by its peak firing rate during normal saccades.

### Modeling of blink-triggered movements

Our objective in this study was to test whether a linear model that took into account saccade and BREM dynamics was sufficient to account for the heterogeneity of blink-triggered movements. If this was indeed true, the saccade could be extracted from linear decomposition of the blink-triggered movement. Previous studies had found that blink-perturbed movements could not be explained as a simple linear combination of saccades and BREMs (Goossens and Van Opstal 2000a). Hence, we created a modified linear model – linear combination with delay. We chose velocity space to model the movements because of the sharper array of features (e.g., number of peaks) in the velocity signal compared to the position signal that could potentially be exploited to fit the movements. The model is described by the following equation:

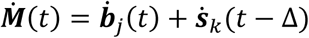

where ***M***(*t*), ***b***(*t*), and ***s***(*t*) represent the position signals of the simulated blink-triggered movement, BREM, and normal saccade, respectively, and the overdot denotes the derivative of those signals, representing velocity. Note that each of the terms is a two-element vector containing values from horizontal and vertical channels. For each of the three movements, we included additional 20 ms snippets on either side of the onset and offset times determined by velocity criteria.

Next, we minimized Euclidean distance in velocity space to fit the simulated movements to recorded blink-triggered movements. The free parameters of the model are *j*, *k*, and Δ, which are the indices of individual BREMs, saccades, and a time delay. For each blink-triggered movement ***m***_*i*_(*t*), we determined the optimal parameters as

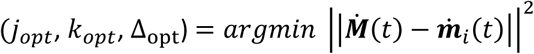

The parameter space was chosen to be computationally efficient and spanning reasonable values. Thus, we randomly chose 100 trials to provide the saccade profiles, 50 trials to provide BREM profiles, and scanned integer time delays in the interval [-100, 100]. Negative values for Δopt indicate that saccade onset precedes BREM onset in the blink-triggered movement, while positive values indicate that saccade onset follows BREM onset. This approach is illustrated in Figure 2a.

**Figure 2.**
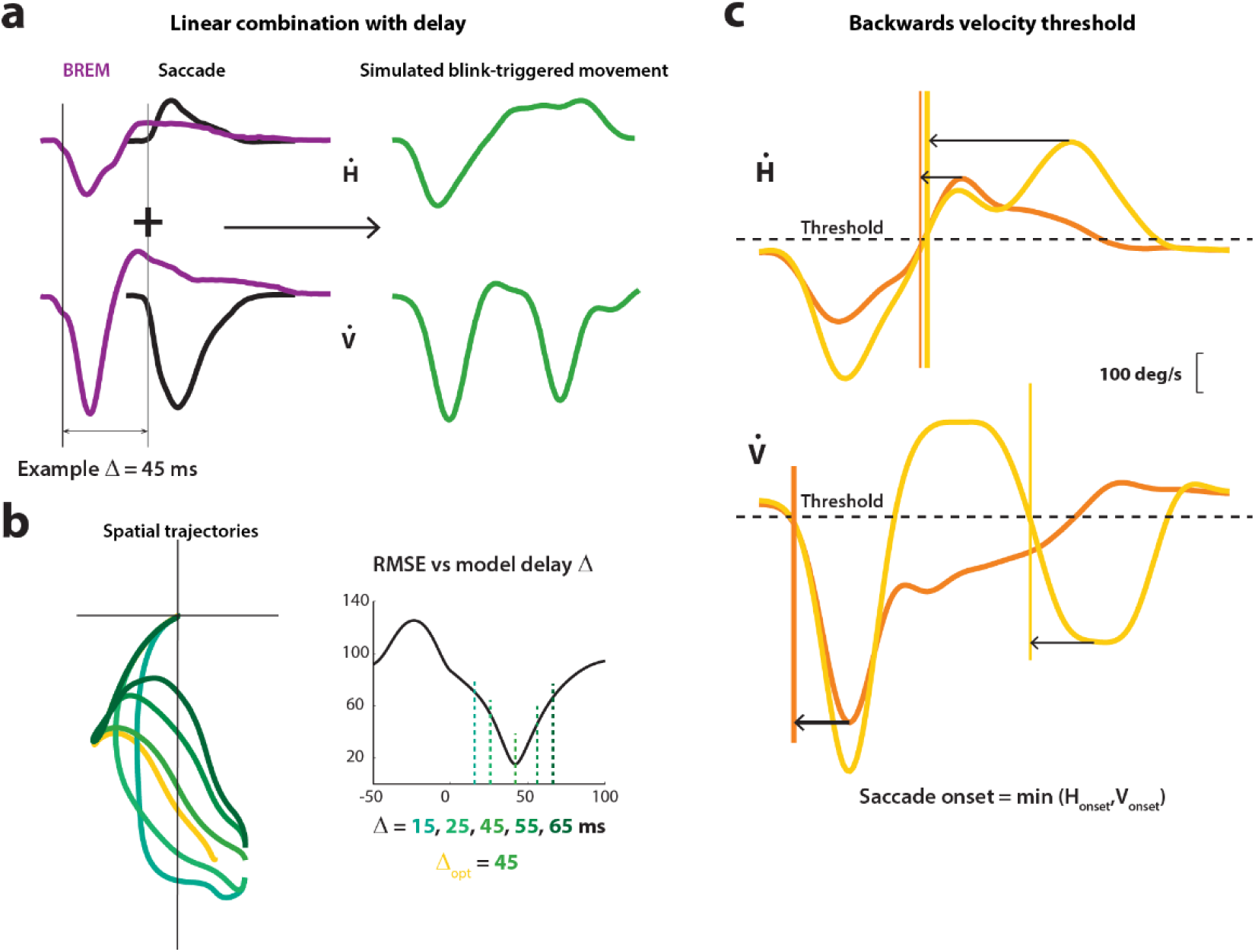
**a**. Schematic of the linear combination approach. The velocity profiles of BREMs (purple traces) and normal saccades (black traces) are shifted in time (e.g, by 45 ms) and added to give a simulated blink-triggered movement (green traces). **b.** Spatial trajectories of an example blink-triggered movement (yellow trace) and five simulated movements with various time shifts (shades of green) are shown on the left. On the right is a plot of the root mean-squared error between various simulated movement profiles and the actual profile as a function of simulated time shift Δ. The value of the time shift that provides the best fit to the actual blink-triggered movement (in this example, the best fit occurs at delay = 45 ms) is considered to be the starting point of the saccade in the combined movement. **c**. Schematic of the backwards threshold approach. Two example blink-triggered movement traces (yellow and orange) from Figure 1 are shown. The time at which velocity crosses a pre-determined threshold (horizontal dotted line), computed backwards from the last peak, is considered as the onset time for that component. The minimum of horizontal and vertical onset times is taken to be the time of saccade onset.

### Detection of back-thresholded saccades

For comparison with the linear combination with delay model, we also detected saccade onset in blink-triggered movements using a previously used backwards velocity threshold method (Gandhi and Bonadonna 2005). In order to do this, we detected the peak in the velocity profile of the blink-triggered movement and marched backwards in time until the standard onset velocity criterion (50 deg/s) was crossed for at least 5 consecutive time points. The idea behind this approach is that the peak velocity of the movement should occur during the saccade component, and going backwards in time until the threshold is crossed should isolate the saccade alone. In several sessions, peak velocity was attained during the initial phase of the blink-triggered movement, which was clearly contributed by the BREM. Hence, we applied the backwards threshold method starting from the second significant peak for these movements. This approach is depicted in Figure 2b.

### Other analyses and statistical tests

For the analyses that involve studying the effect on model performance of an external parameter (e.g., direction or optimal delay), we binned trials from all sessions according to that parameter, and computed the average mean-squared error (MSE) across trials for each bin. Significant trends were identified by comparing these to the null distribution (uniform distribution/no trend) generated by a bootstrap approach with appropriate confidence intervals that took into account the number of trials available in a given bin. Where applicable, we used the Wilcoxon-rank-sum test for a two-way comparison of distribution medians.

In addition to MSE, we also used Spearman’s rank correlation to compare the efficiency of different approaches in generating velocity profiles that closely match the control profiles. A point-by-point correlation of two profiles ignores magnitude differences (that is represented in the MSE) and tests whether their shapes are similar. Scaled profiles would result in a correlation of 1.

We also computed a suppression index (SI) for individual neurons as the relative change in the activity for extracted saccade trials with respect to control trials. We computed the index using the average activity in a 40 ms window around saccade onset as,

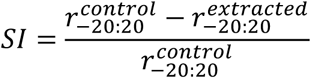

where 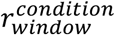 is the average activity in *window* around saccade onset for *condition* trials.

## Results

In order to extract saccades from blink-triggered movements, we needed three sets of data. In animals performing the delayed saccade task, during each session, we recorded 1) a set of normal saccades to two or more targets (typically more than 100 per session), 2) a set of blink-triggered gaze shifts to those targets by inducing reflex blinks after the go cue (around 100 trials per session), 3) blink-related eye movement (BREM) profiles by evoking blinks during initial fixation, before peripheral target onset (around 50 per session). We had 100 session-target pairs in all, and analyzed data for each session-target pair separately.

First, we simulated a range of movement profiles by linear summation of randomly chosen saccade and BREM velocity profiles at all possible time shifts relative to each other. Then, for each real blink-triggered movement, we determined the simulated profile that best fit it (see Methods), and the associated triad of saccade, BREM and time shift parametrized the decomposition of the blink-triggered movement. Note that the optimal time shift effectively determined the time of saccade onset within the movement. Figure 3a shows the average blink-triggered movement velocity profiles for one session-target pair (orange traces) and the corresponding best-fitting simulated movement profile average (green traces). Next, we extracted the underlying saccade by subtracting the respective BREM profile from each blink-triggered movement at the appropriate time shift. The dynamics of the extracted saccades closely resembled normal saccades, as judged from the shape of their velocity profiles (Figure 3b, compare blue vs black traces). For comparison, we also computed an alternative saccade onset time using a backwards velocity threshold criterion (see Methods) that has been used previously (Gandhi and Bonadonna 2005). Note that the dynamics of the saccade estimated using this criterion (Figure 3b, red traces) deviated significantly from that of normal saccades or those extracted using the aforementioned method.

**Figure 3.**
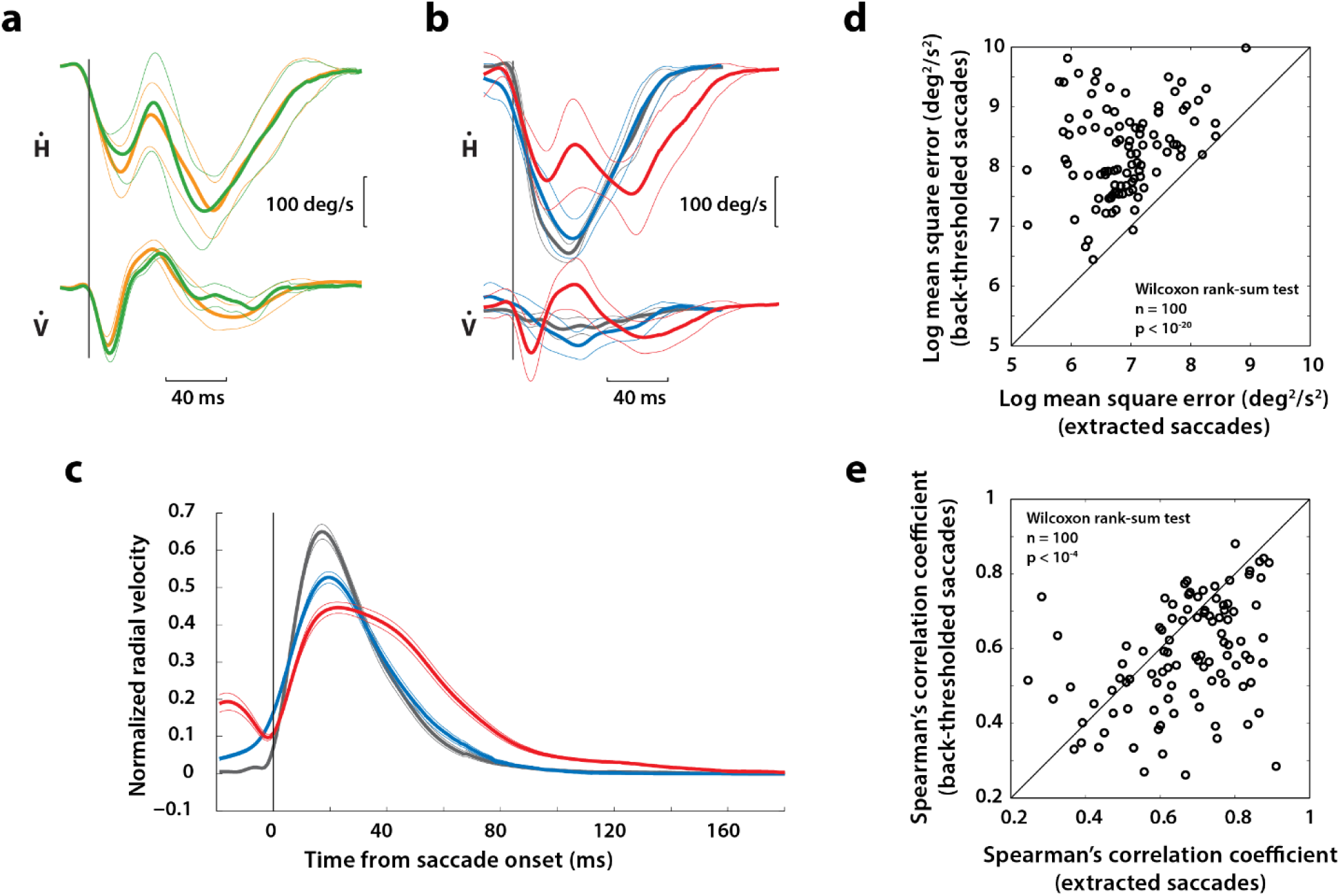
**a**. Average horizontal and vertical velocity profiles of actual (orange traces) and simulated best-fitting (green traces) blink-triggered movements for an example session-target pair. The thin lines are the standard error bounds across trials. **b**. Average horizontal and vertical velocity profiles of normal saccades (black), extracted saccades (blue), and back-thresholded saccades (red) for the same session-target pair as in **a**. **c**. Average normalized radial velocity profiles across all sessions for the three saccade types shown in **b**. **d**. Log-log plot of the mean-squared error between extracted and normal saccade velocity profiles (abscissa) plotted against back-thresholded and normal velocity profiles (red bars) across session-target pairs. Each point is from a session-target pair. **e**. Spearman’s rank correlation between extracted and normal saccade velocity profiles plotted against the correlation between back-thresholded and normal velocity profiles.

Figure 3c shows the normalized, average radial velocity profiles across all session-target pairs for normal, extracted and back-thresholded saccades. Qualitatively, the extracted saccades overlap substantially with normal saccades. Some of the discrepancies could be because the dynamics of the motor command, and therefore the saccade, triggered in response to a blink may be slightly attenuated (for more, see below). Moreover, the velocity profiles of extracted saccades were much closer to normal saccades than those of back-thresholded saccades. To quantify this observation, we computed the mean square error (MSE, equivalently, Euclidean distance in velocity phase space) between normal and extracted saccades, as well as between normal and back-thresholded saccades, for each session-target pair. Figure 3d shows the MSEs for the two cases plotted against each other (blue – normal vs extracted, red – normal vs back-thresholded). The MSEs were significantly lower for extracted saccades compared to back-thresholded saccades (Wilcoxon rank-sum test, n=100, p < 10^-20^). Note that the MSE reflects differences between two profiles regardless of their shape or where the deviations occur. To further evaluate whether the shape of the extracted saccade profiles resembled that of normal saccades, we computed the correlation between the velocity traces for each pair considered above. Two traces with similar shapes (e.g., scaled versions of each other) should have a high correlation coefficient compared to traces of dissimilar shapes. Figure 3e shows a scatter plot of the session-wise correlation values (Spearman’s correlation coefficient) for the two pairs of traces (normal vs extracted, mean +/- s.d. = 0.71 +/- 0.14; normal vs back-thresholded, mean +/- s.d. = 0.6 +/- 0.13). The correlations were significantly higher for extracted saccades (Wilcoxon rank-sum test, n=100, p < 10^-4^). Taken together, these results suggest that extraction of saccades from blink-triggered movements by linear decomposition is superior to the currently used method of determining the saccadic component and is likely more veridical in its estimates of saccade onset and dynamics.

Since the BREMs are stereotypical in a given animal and always excurse in roughly the same direction from the point of fixation, it is reasonable to expect an approach like ours to have differential efficacy depending on direction of the intended saccade. We therefore wanted to verify whether the performance of the algorithm was equally good for blink-triggered movements to all directions. We binned trials across all sessions into different directions based on the direction of the actual movement (i.e., final eye position relative to initial eye position) and computed the average MSE in each bin. This is shown in Figure 4a (thick blue trace; thin traces represent +/- s.e.m.). For comparison with the null (uniform) profile, we computed the bootstrapped version of this trace by randomly assigning trials to direction bins (thin cyan represent mean and +/- 95% confidence intervals). The MSEs were largely uniform across directions, expect for a slight increase for movements to the bottom right quadrant, and a slight decrease for movements to the bottom left quadrant. Intriguingly, the BREMs were always directed to the bottom left quadrant for all our subjects. We also performed this analysis for the backwards threshold approach, as shown in Figure 4b (red curves; magenta curves show the bootstrapped distribution). The MSEs were more asymmetric and dependent on direction for back-thresholded saccades, suggesting that a simple threshold criterion may ignore the complex manner in which BREMs and saccades may interact dependent on direction.

**Figure 4.**
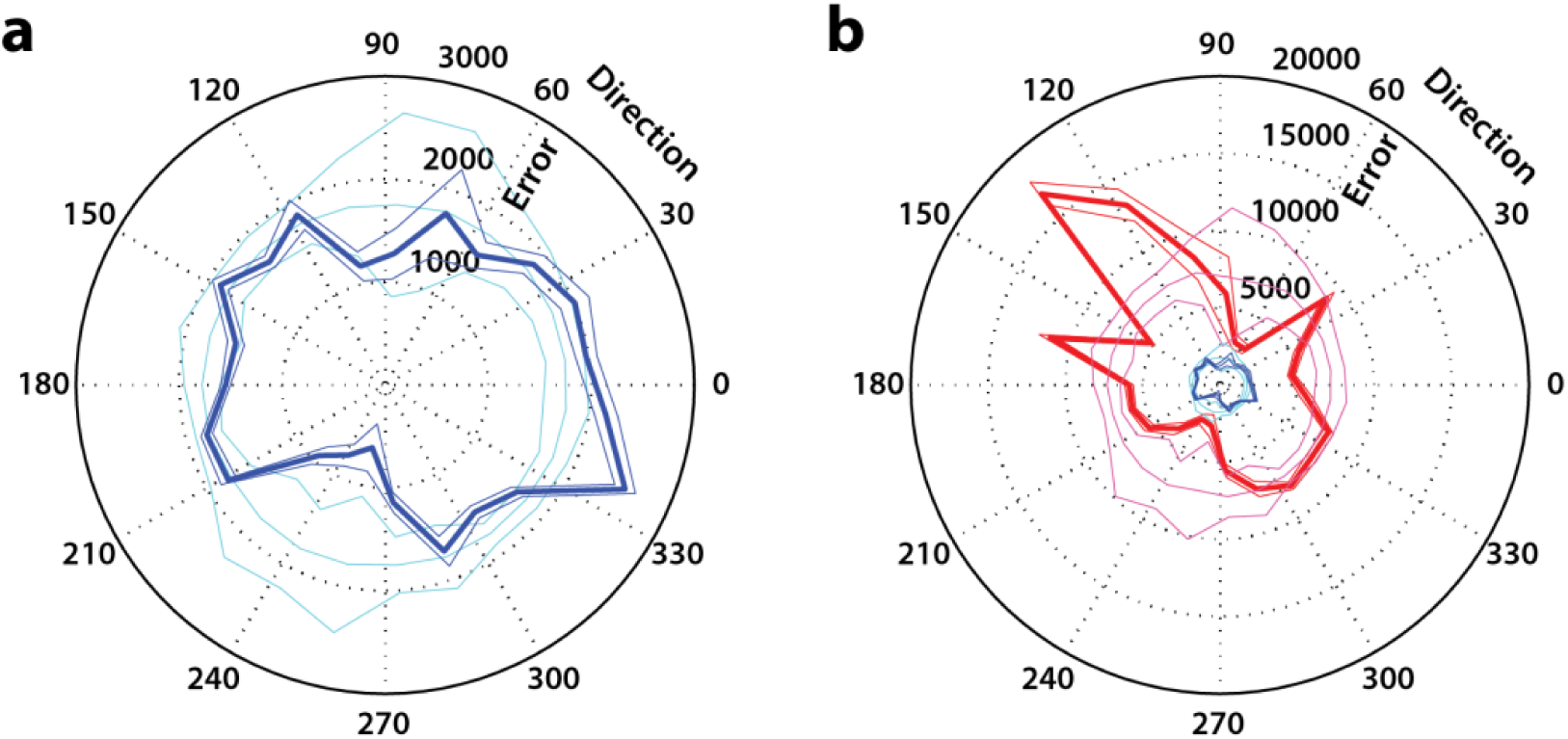
**a**. Polar plot of the mean-squared error between individual extracted saccades and the average control saccade as a function of movement direction (blue traces). The thick blue trace is the average error across trials in a given direction bin, and the thin traces are +/- s.e.m. The cyan traces represent the average bootstrapped error and +/- 95% CI, and provide the null distribution in a given direction for comparison. **b.** Same as **a**, but for back-thresholded saccades (red traces), and the corresponding bootstrapped error distribution (magenta traces). The plot from **a** is also shown (blue traces near the centre) for comparison of magnitude.

It is pertinent to remark here that the method of linear combination is certain to provide an optimal decomposition of a blink-triggered movement, since any set of simulated movements is going to contain one that fits the actual movement best. Although the results presented above demonstrate the method’s utility in extracting saccades with near-normal velocity profiles, its use in studies of the neural mechanisms of movement preparation requires evidence that saccade-related neural activity is not unduly affected by the re-computed onset times. In order to determine whether this was true, we compared motor bursts of 51 neurons in the superior colliculus (SC) for normal saccades, and both extracted and back-thresholded saccades from blink-triggered movements. We only included session-target pairs in which the target was in the response field of the recorded SC neuron. Figure 5a shows the trial-averaged spike density of one neuron, aligned on respective saccade onsets, for each of the three cases. Note the remarkable similarity between the burst profiles for normal and extracted saccades (blue vs black traces). Figure 5b shows the average population activity for the three cases. To quantify the similarity of the bursts, we performed the correlation analysis presented earlier for the velocity profiles in Figure 3e. The distribution of correlation coefficients (Spearman’s rank correlation) for the two pairs of comparisons plotted against each other is shown in Figure 5c. Correlations were significantly higher for the extracted saccade bursts (Wilcoxon rank-sum test, n=51, p < 10^-6^), suggesting that linear decomposition with delay largely preserves the saccade-related burst compared to back-thresholding.

**Figure 5.**
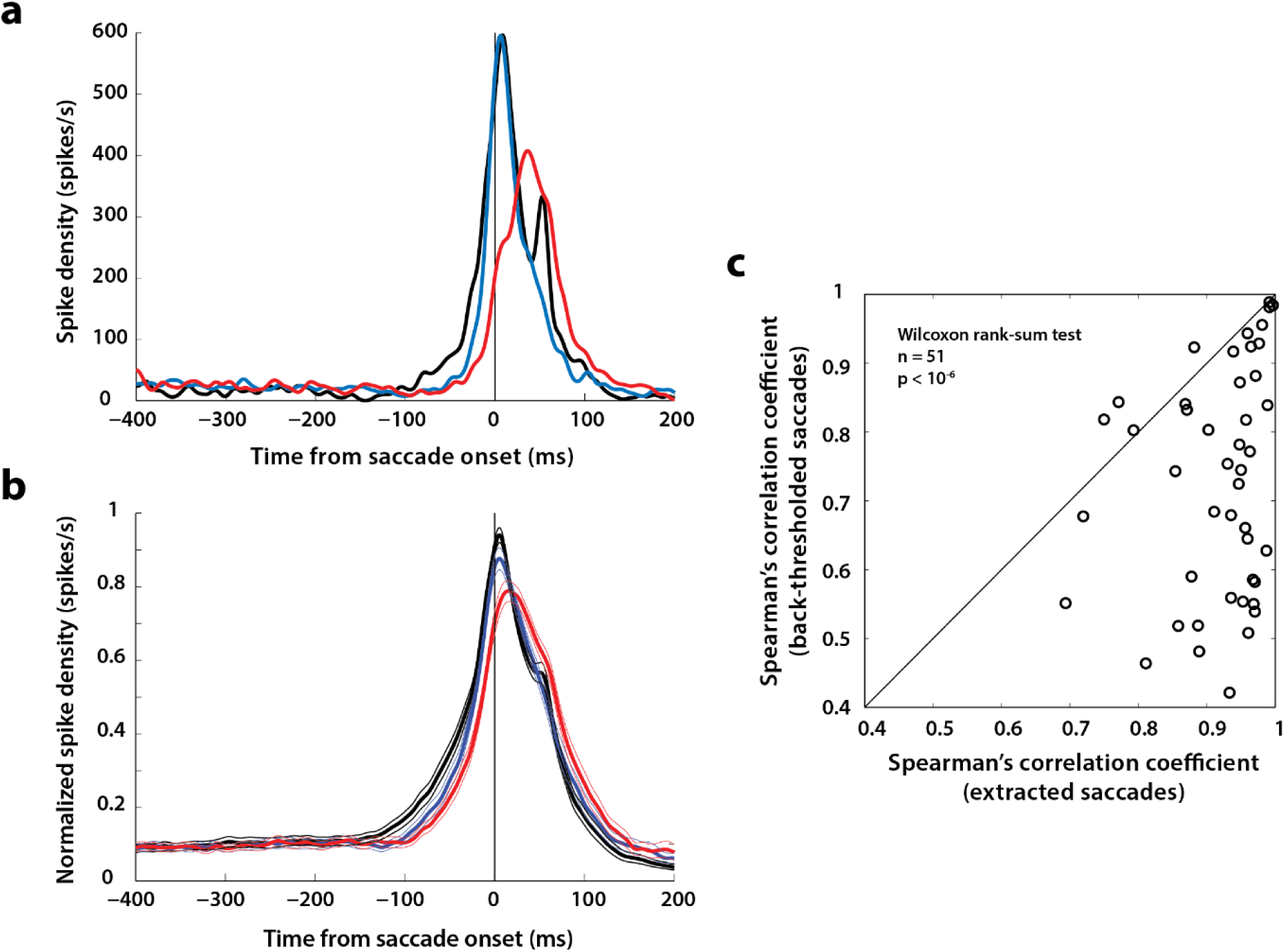
**a**. Saccade-related burst for one example SC neuron for normal (black), extracted (blue), and back-thresholded (red) saccades. **b**. Population-averaged motor burst in SC for the three types of saccades in **a**. **c**. Spearman’s rank correlation between extracted and normal motor bursts plotted against the correlation between back-thresholded and normal motor bursts. Each point corresponds to one neuron.

Previous studies have observed strong attenuation in the saccadic burst in a few SC neurons (Goossens and Van Opstal 2000b), but no suppression is apparent in our data (Figure 5a, b). To resolve this discrepancy, we more closely analyzed the timing of the saccades extracted using our approach relative to the occurrence of the blink. Figure 6a shows the distribution of extracted saccade onset times relative to overall movement onset across all combined blink-saccade movements. In other words, this is the distribution of optimal time shifts at which linearly combining a BREM and saccade would produce each blink-triggered movement. Saccade onset followed BREM onset in the majority of trials. Observe that saccades can begin as late as 80 ms into the movement – an eon in the time scale of sensorimotor integration - highlighting the importance of precisely determining saccade onset time for studying movement preparation using the blink approach. Intriguingly, the distribution of optimal delays for saccade onset was bimodal, with an early peak around 10 ms and a later peak around 40 ms after movement onset. We thought this might provide a clue to the issue of suppression. Note that when saccade onset follows the blink by as early as 0-20 ms, the blink is occurring at a time when the command to generate a saccade has already been issued (accounting for efferent delays), and is likely to affect the burst when it is approaching peak value. If so, any suppression in these cases will alter the dynamics of the underlying programmed saccade and affect the ability of the algorithm to extract a control-like saccade.

**Figure 6.**
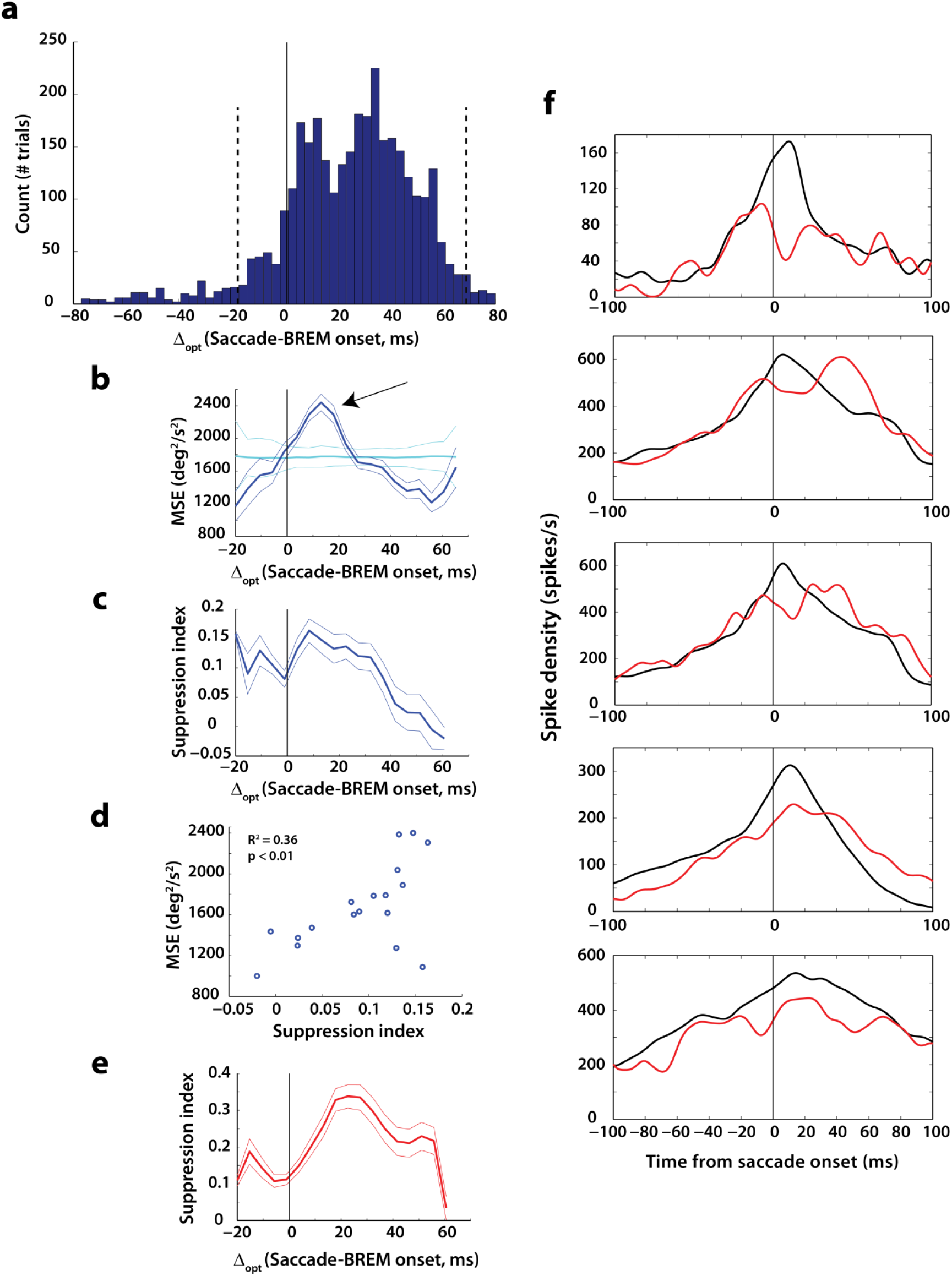
**a**. Distribution of extracted saccade onset times relative to overall movement onset (delay, Δopt) for all trials. Based on the method, saccade onset occurred after movement onset (or, equivalently, BREM onset) in the majority of trials. The vertical dotted lines represent the 95^th^ percentile of the distribution and the analyses were restricted to trials in this window of delays. **b**. Average mean-squared error between extracted and control saccade velocity profiles as a function of optimal delay (thick blue trace) and +/- s.e.m (thin blue traces). The cyan traces show the bootstrapped null distribution and +/- 95% CI for a test of uniformity. Arrow indicates region where MSE is greater than null distribution (see text for details). **c.** Neural suppression index (relative suppression for extracted saccades compared to control saccades) as a function of optimal delay. **d.** Correlation between mean-squared error and suppression index (points in **b** vs **c**). Each point is from a bin of optimal delays. **e.** Neural suppression index for back-thresholded saccades as a function of optimal delay, for better comparison with previous results. Note that relative suppression is higher when aligned with respect to the back-thresholded saccade, compared to that in **c**. **f.** Examples of suppression in five different neurons. Black traces show the trial-averaged activity aligned to saccade onset in control trials, and red traces depict the activity aligned to back-thresholded saccades, averaged across trials with optimal delays where suppression is highest (10-30 ms from **e**).

We first tested whether the performance of the linear combination with delay approach was a function of the optimal delay identified by the algorithm (Figure 6b). The MSE was indeed higher for optimal delays between 0-20 ms (blue traces), indicated by the increase above the bootstrapped baseline (cyan traces). We then computed the suppression index (see Methods) for each neuron as a function of optimal delay bin. The average index across the population is shown in Figure 6c (thick trace; thin traces represent +/- s.e.m.). Note that the suppression is highest in the 0-20 ms window. Indeed, neuronal suppression was a reliable predictor of the MSE, as seen in the correlation of the two variables across bins (Figure 6d, R^2^ = 0.36, p < 0.01). However, the magnitude of suppression was still lower (15% at its maximum in Figure 6c) than previously observed (Goossens and Van Opstal 2000b). It is important to note that the suppression values so far were calculated based on aligning activity to extracted saccade onset, which seeks to maximize the similarity of the underlying saccade to a control movement (and may thus minimize differences at the neural level as well), whereas, in previous studies, activity was aligned to saccade onset estimated using the backwards velocity threshold criterion. Suppression magnitudes were indeed higher for back-thresholded saccades, peaking at 35% at the population level (Figure 6e). Finally, we wanted to verify whether individual neurons exhibited suppression that resembled the strong attenuation observed in the previous study. Figure 6f shows the average saccade-aligned burst profiles for five individual neurons for control (black traces) and back-thresholded saccades (red traces), in trials that fall within the 0-30 ms window of optimal delays estimated using the linear combination approach. In all five examples, a strong reduction/dip in the activity is evident during the peri-saccadic period. Thus, it is possible that the strong suppression observed in the previous study in a handful of neurons may be a result of a combination of factors, including timing of the saccade relative to the blink.

## Discussion

Current perspective holds that rapid redirection of the visual axis during a blink cannot be accounted for by a sum of a typical saccade (Goossens and Van Opstal 2000a) and a BREM and that blinks are associated with potent suppression of activity in premotor circuitry of the saccadic system (Goossens and Van Opstal 2000b). In contrast, we have shown that blink-triggered movements can be characterized as a linear combination of BREMs and saccades with an arbitrary delay that can be estimated from the movement. Furthermore, we found that the dynamics of saccades extracted using this method as well as the associated motor bursts in SC are similar to those of normal saccades.

The discrepancies between our results and the previous studies could be due to several factors. First, in the previous behavioral study, the possibility of a staggered, time-shifted linear superposition was not systematically explored (Goossens and Van Opstal 2000a). As we show here, BREM onset and saccade onset can be offset by several tens of milliseconds, and taking this time shift into account can increase the explanatory power of the linear analysis significantly. Second, we show that the saccade-related burst is largely preserved for the extracted saccade. We believe this discrepancy could be due to the fact that our analyses included a large portion of blink-*triggered* saccades, i.e., movements in which blink onset preceded saccade onset by several tens of milliseconds, whereas previous studies focused largely on co-occurring blinks and saccades. Since our laboratory’s focus is to use the reflex blink as a tool to probe underlying motor preparation, we induced blinks early, and thus obtained many trials where the blink likely triggered neural processes that resulted in a saccade much later. In the previous studies, blinks were timed to occur during typical saccade reaction times, likely resulting in instances where the saccadic burst had commenced and was perturbed by the blink. This notion is consistent with the observation that the suppression on blink-perturbed trials, especially for saccade onset times computed using previous approaches, is maximal around 20 ms after the onset of the BREM, which is when the peak of the burst would have occurred if the final motor command was issued at the time of BREM (Figure 6e). In other words, we think the likelihood of observing suppression is higher when the blink interferes with the motor burst around the time of its peak. These claims are further bolstered by the observation that the algorithm fails to extract saccades resembling control saccades for this window of optimal delays (higher MSE, arrow in Figure 6b), possibly due to altered kinematics as a result of the attenuated burst peak and by the strong suppression of the peak in example individual neurons for trials that fall in this window of delays (Figure 6f). For true blink-triggered saccades, i.e., for longer optimal delays between blink and saccade onset, suppression of low-frequency preparatory activity, if any, may have enough time to recover to produce a full-fledged saccadic burst. The dynamics of saccades extracted from such blink-triggered movements do not seem to be very different from those of normal saccades; thus it is not surprising that SC activity for the two types of saccades are similar. It should be noted here that, even disregarding the specific focus on the time course of the saccade relative to the blink, the intensity of the motor burst was attenuated when we used the velocity-based threshold criterion (Figure 5b).

Another possible explanation for the discrepancy in the extent of attenuation is differences in the air puff stimuli used to induce reflex blinks. The observed short latency attenuation of saccade-related activity of a small subset of neurons in the previous study was locked to the time of air puff delivery, and as such, may have been linked to the strength of the air puff. It is entirely possible that the air puffs we used were weaker, and combined with early delivery times (when preparatory activity is lower), precluded the strong attenuation described previously. It should further be noted that the previous study observed strong suppression in only a small percentage of the overall population, and we observed a similar strong effect in several individual neurons (some of which are shown in Figure 6f). The primary purpose of this study was to find a way to precisely determine saccade onset from a blink-triggered movement, and we think a systematic analysis of these multiple factors and a thorough re-verification of previous results is beyond its scope.

The fact that BREMs and saccades can be linearly combined to produce the observed movement suggests that the underlying neural processes producing a BREM and saccade are independent. Although the neural correlates of a BREM are unclear, we have previously shown that the BREM itself does not affect activity in caudal SC (Jagadisan and Gandhi 2016). However, the BREM has been shown to be associated with a pause in the activity of OPNs (Schultz et al. 2010), which we think allows activity in premotor structures such as SC to activate the burst generators and produce a saccade (Gandhi and Bonadonna 2005), while the BREM itself reaches completion in parallel. We have exploited the observation that blinks remove a potent source of inhibition on the oculomotor system to study latent dynamics of movement cancellation (Walton and Gandhi 2006), characteristics of SC stimulation-evoked saccades under extended disinhibition (Katnani et al. 2012), and the relationship between target selection and motor preparation (Katnani and Gandhi 2013). In addition to the BREM and saccade components of blink-triggered movements, leakage of premotor activity before saccade onset (but after blink onset) may produce slow eye movements, as seen in Figure 2c (non-zero velocity before t=0 in the blue trace). Others have used blinks to study initiation interactions not just for saccades, but also smooth pursuit (Rambold et al. 2004) and vergence (Rambold et al. 2002). Collectively, these observations, along with the demonstration of linear independence, suggest that the reflex blink may be a simple and elegant tool to probe the dynamics of movement preparation and execution in the saccadic system.

## Notes

**Conflict of interest**: The authors declare no competing financial interests.

## References

Bryant CL, and Gandhi NJ. Real-time data acquisition and control system for the measurement of motor and neural data. J Neurosci Methods 142: 193–200, 2005.

Cohen B, and Henn V. Unit activity in the pontine reticular formation associated with eye movements. Brain Res 46: 403–410, 1972.

Gandhi NJ, and Bonadonna DK. Temporal interactions of air-puff evoked blinks and saccadic eye movements: insights into motor preparation. J Neurophysiol 93: 1718–1729, 2005.

Gandhi NJ, and Katnani HA. Motor functions of the superior colliculus. Annu Rev Neurosci 34: 203–229, 2011.

Goossens HH, and Van Opstal AJ. Blink-perturbed saccades in monkey. I. Behavioral analysis. J Neurophysiol 83: 3411-3429, 2000a.

Goossens HH, and Van Opstal AJ. Blink-perturbed saccades in monkey. II. Superior colliculus activity. J Neurophysiol 83: 3430-3452, 2000b.

Jagadisan UK, and Gandhi NJ. Disruption of fixation reveals latent sensorimotor processes in the superior colliculus. J Neurosci 36: 6129–6140, 2016.

Katnani HA, and Gandhi NJ. Time course of motor preparation during visual search with flexible stimulus-response association. J Neurosci 33: 10057–10065, 2013.

Katnani HA, Van Opstal AJ, and Gandhi NJ. Blink perturbation effects on saccades evoked by microstimulation of the superior colliculus. PLoS One 7: e51843, 2012.

Keller EL. Participation of medial pontine reticular formation in eye movement generation in monkey. J Neurophysiol 37: 316–332, 1974.

Keller EL, Gandhi NJ, and Shieh JM. Endpoint accuracy in saccades interrupted by stimulation in the omnipause region in monkey. Vis Neurosci 13: 1059–1067, 1996.

Krauzlis RJ, Lovejoy LP, and Zenon A. Superior colliculus and visual spatial attention. Annu Rev Neurosci 36: 165–182, 2013.

Luschei ES, and Fuchs AF. Activity of brain stem neurons during eye movements of alert monkeys. J Neurophysiol 35: 445–461, 1972.

Rambold H, El Baz I, and Helmchen C. Differential effects of blinks on horizontal saccade and smooth pursuit initiation in humans. Exp Brain Res 156: 314–324, 2004.

Rambold H, Sprenger A, and Helmchen C. Effects of voluntary blinks on saccades, vergence eye movements, and saccade-vergence interactions in humans. J Neurophysiol 88: 1220–1233, 2002.

Rottach KG, Das VE, Wohlgemuth W, Zivotofsky AZ, and Leigh RJ. Properties of horizontal saccades accompanied by blinks. J Neurophysiol 79: 2895–2902, 1998.

Schultz KP, Williams CR, and Busettini C. Macaque pontine omnipause neurons play no direct role in the generation of eye blinks. J Neurophysiol 103: 2255–2274, 2010.

Walton MM, and Gandhi NJ. Behavioral evaluation of movement cancellation. J Neurophysiol 96: 2011–2024, 2006.

